# K_v_7 Channel Opener Retigabine Reduces Self-Administration of Cocaine but Not Sucrose in Rats

**DOI:** 10.1101/2023.05.18.541208

**Authors:** Esteban S. Urena, Cody C. Diezel, Mauricio Serna, Grace Hala’ufia, Lisa Majuta, Kara R. Barber, Todd W. Vanderah, Arthur C. Riegel

## Abstract

The increasing rates of drug misuse highlight the urgency of identifying improved therapeutics for treatment. Most drug-seeking behaviors that can be modeled in rodents utilize the repeated intravenous self-administration (SA) of drugs. Recent studies examining the mesolimbic pathway suggest that K_v_7/KCNQ channels may contribute in the transition from recreational to chronic drug use. However, to date, all such studies used noncontingent, experimenter-delivered drug model systems, and the extent to which this effect generalizes to rats trained to self-administer drug is not known. Here, we tested the ability of retigabine (ezogabine), a K_v_7 channel opener, to regulate instrumental behavior in male Sprague Dawley rats. We first validated the ability of retigabine to target experimenter-delivered cocaine in a CPP assay and found that retigabine reduced the acquisition of place preference. Next, we trained rats for cocaine-SA under a fixed-ratio or progressive-ratio reinforcement schedule and found that retigabine-pretreatment attenuated the self-administration of low to moderate doses of cocaine. This was not observed in parallel experiments, with rats self-administering sucrose, a natural reward. Compared to sucrose-SA, cocaine-SA was associated with reductions in the expression of the K_v_7.5 subunit in the nucleus accumbens, without alterations in K_v_7.2 and K_v_7.3. Therefore, these studies reveal a reward specific reduction in SA behavior considered relevant for the study of long-term compulsive-like behavior and supports the notion that K_v_7 is a potential therapeutic target for human psychiatric diseases with dysfunctional reward circuitry.

## INTRODUCTION

Repeated exposure to addictive drugs is known to hyperactivate the mesocorticolimbic ventral tegmental area (VTA), nucleus accumbens (NAc), and prefrontal cortex (PFC) brain regions important for the integration of reinforcing stimuli with goal-directed behavior ^1,2^. A promising approach to counteract this integration is stabilization of K_v_7/KCNQ voltage-gated potassium channels, which express different heteromeric combinations of K_v_7.2/K_v_7.3, and K_v_7.3/K_v_7.5 subunits that strongly influence channel properties ^3–8^. In neurons, the opening of K_v_7 channels triggers a repolarizing M-current associated with spike frequency adaptation [9,10]. Retigabine, a K_v_7 channel opener, stabilizes M-currents and by positive modulation of the membrane potential restricts the excitability of cortical and mesencephalic dopamine (DA) neurons ^9,10^. As such, targeting K_v_7 channels with retigabine may prove useful in modulating dysfunctional circuitry associated with reward-related neuropsychiatric and other neurological diseases ^11–13^.

Furthermore, *in vitro* and *in vivo* studies show that retigabine can reduce the release of dopamine from the terminals of mesencephalic dopaminergic neurons ^14^ and reduce the locomotor activity induced by psychostimulants ^15,16^. These actions of retigabine are reversed by the K_v_7 channel blocker XE-991 ^15,17^. However, to date, most studies using retigabine to modulate drug-induced changes in dopamine neurons have used noncontingent methods ^16,18^. Our previous work used an operant model to show that during re-exposure to drug-predictive cues, retigabine reduced cocaine-seeking behavior and the related changes in pyramidal cell firing and K_v_7 channel currents ^9^. However, it remains unknown whether retigabine alters ongoing drug SA behavior, from which these relapse-like neuroadaptations later emerge.

In this study, we propose that K_v_7 channels act as inhibitory regulators of motivated drug behavior, thus decreasing the reinforcing actions of cocaine during the maintenance of chronic voluntary self-administration. We first tested this hypothesis in male Sprague Dawley rats to identify a retigabine dose sufficient to reduce the conditioned preference for cocaine and then determined whether lower, similar or higher doses of retigabine reversibly reduced operant responding for various doses of cocaine. We showed that retigabine reversibly weakens the motivation for cocaine-SA at low to moderate doses on a fixed-ratio or progressive-ratio reinforcement schedule. We examined the expression of various K_v_7 channel subunits in relevant mesolimbic regions and provided evidence that relative to sucrose-SA, chronic cocaine-SA was associated with a notable reduction in the expression of the K_v_7.5 channel subunit in the accumbens. This study shows that the motivation to self-administer cocaine is reduced by a potent activator of K_v_7 channels and that the ability to ameliorate this drug behavior does not necessarily generalize to other nondrug (natural) rewards.

## METHODS

### Subjects

Male Sprague Dawley rats (n=120; Envigo) weighing 250-275g were housed individually in a reverse 12-hour light / dark cycle (light off at 06:00 am). Rats were divided into two groups: fed ad libitum and food restricted. Ad libitum rats had non-restricted access to food and water; food-restricted rats had free access to water and were food restricted to maintain 80% of the body weight of ad libitum rats. All procedures performed were preapproved by the Institutional Animal Care and Use Committee of the University of Arizona and according to the Animal Care Guidelines of the National Institutes of Health for the Care and Use of Laboratory Animals.

### Conditioned place preference (CPP)

Using our established procedures ^19^, we performed cocaine CPP in a three-chambered system (San Diego Instruments). Using a biased design, rats were assigned to the least preferred chamber for pairing with cocaine (10 mg/kg, i.p.). Conditioning sessions (30 min) occurred twice a day, with one chamber in the morning (session-1) and the opposite chamber in the afternoon (session-2). Daily saline and cocaine sessions were counterbalanced with respect to session (Fig. 1A). Pretreatments with retigabine (5 mg/kg, i.p.) or vehicle (10% tween-80 in saline) occurred 15 minutes before each cocaine conditioning session. After the five conditioning days, the rats underwent a 15 min post-test. CPP score was calculated as the time spent in each chamber during post-test minus the time spent in the same chamber during pre-test.

**Fig. 1.**
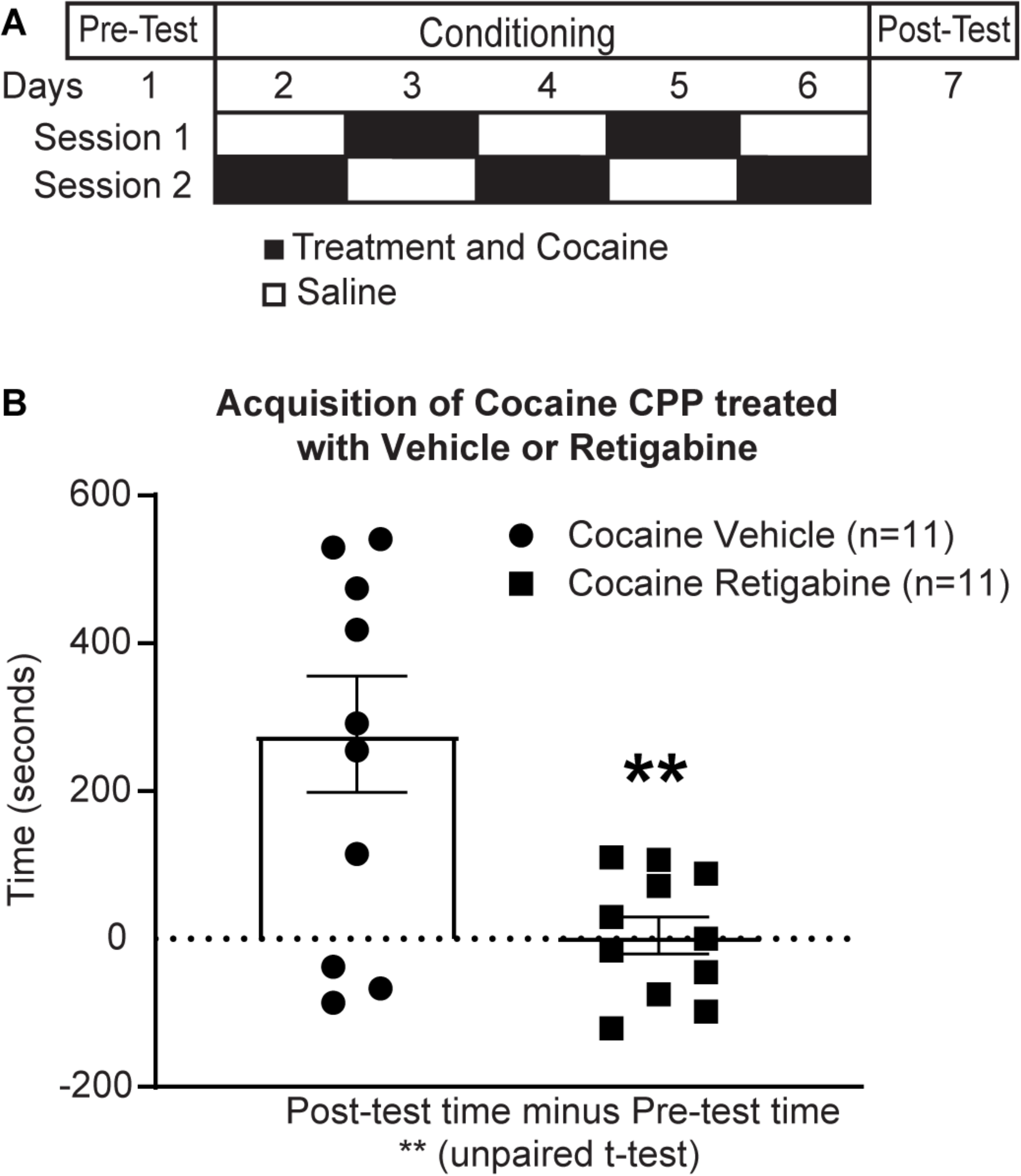
Retigabine reduces the acquisition of cocaine CPP. **A** CPP testing included a pretest and 5 days of twice daily intraperitoneal injections of cocaine (black box; 10 mg/kg) and saline (white box; 1 ml/kg) in alternating counterbalanced fashion. At 15 min before cocaine, rats received intraperitoneal pretreatments of either vehicle (1 ml/kg; n=11) or retigabine (5 mg/kg; n=11). On day 7, rats were tested for side preference. **B** The preference score was calculated as the time spent in each chamber during post-test minus the time spent in the same chamber during pre-test. Cocaine CPP was evident in rats pretreated with vehicle but not retigabine. For these and all other figures, the error bars indicate the mean ± SEM. **p=0.0036 compared to vehicle pretreatment using an unpaired t-test

### Intravenous catheter surgery

Using our published procedures ^9^, rats anesthetized with isoflurane gas (5% for induction; 2.1-2.5% for maintenance) received a 22 gauge silastic catheter placed in the right jugular vein. Catheters were exteriorized to the midscapular region of the back and attached to a vascular access button (Instech, VABR1B/22). After surgery, rats received ketorolac (2 mg/kg, IP) and recovered 4-7 days with daily catheter flushes of a heparinized solution with gentamicin (8mg/kg) (Henry Schein, part # 1098195). During SA-training, catheters were flushed daily with 0.1 ml of 500u/ml heparin dissolved in sterile saline followed by a 0.1 ml lock of heparin (300u/ml).

### Cocaine-SA

Food-restricted rats underwent cocaine-SA (0.5 mg/kg/infusion) on a fixed-ratio 1 (FR1) reinforcement schedule (5-s infusion and 20-s timeout post infusion) for 2h per session, 1 session per day for 6 days per week ^9^. Sessions occurred in sound-attenuating operant chambers connected to Med-PC IV software (Med Associates). During training sessions, a 100μl infusion of cocaine was paired with light and tone cues (white stimulus light above the active lever; 78-dB, 2900-Hz tone). Lever presses on the ‘inactive’ lever were counted but lacked programmed consequences. Seven rats were excluded from the study due to surgical or behavioral complications.

Following acquisition of cocaine-SA, animals were divided into three groups (1A,1B,1C) (**Fig. 2A**). Before retigabine testing, animals in Group-1A,-1B continued training on a FR1 (2h) schedule, at a dose of 0.5 mg/kg/infusion (Group-1A) or a lower dose of 0.1 mg/kg/infusion (Group-1B). Animals were habituated to the (IP) pretreatment injections with a minimum of two days of saline treatments (1 ml/kg, IP) before retigabine and every day thereafter if not receiving retigabine (**Fig. 3A**). The half-life of retigabine in rodents is about 3h ^20^, and we allowed a minimum of 2 days between the retigabine tests.

**Fig. 2.**
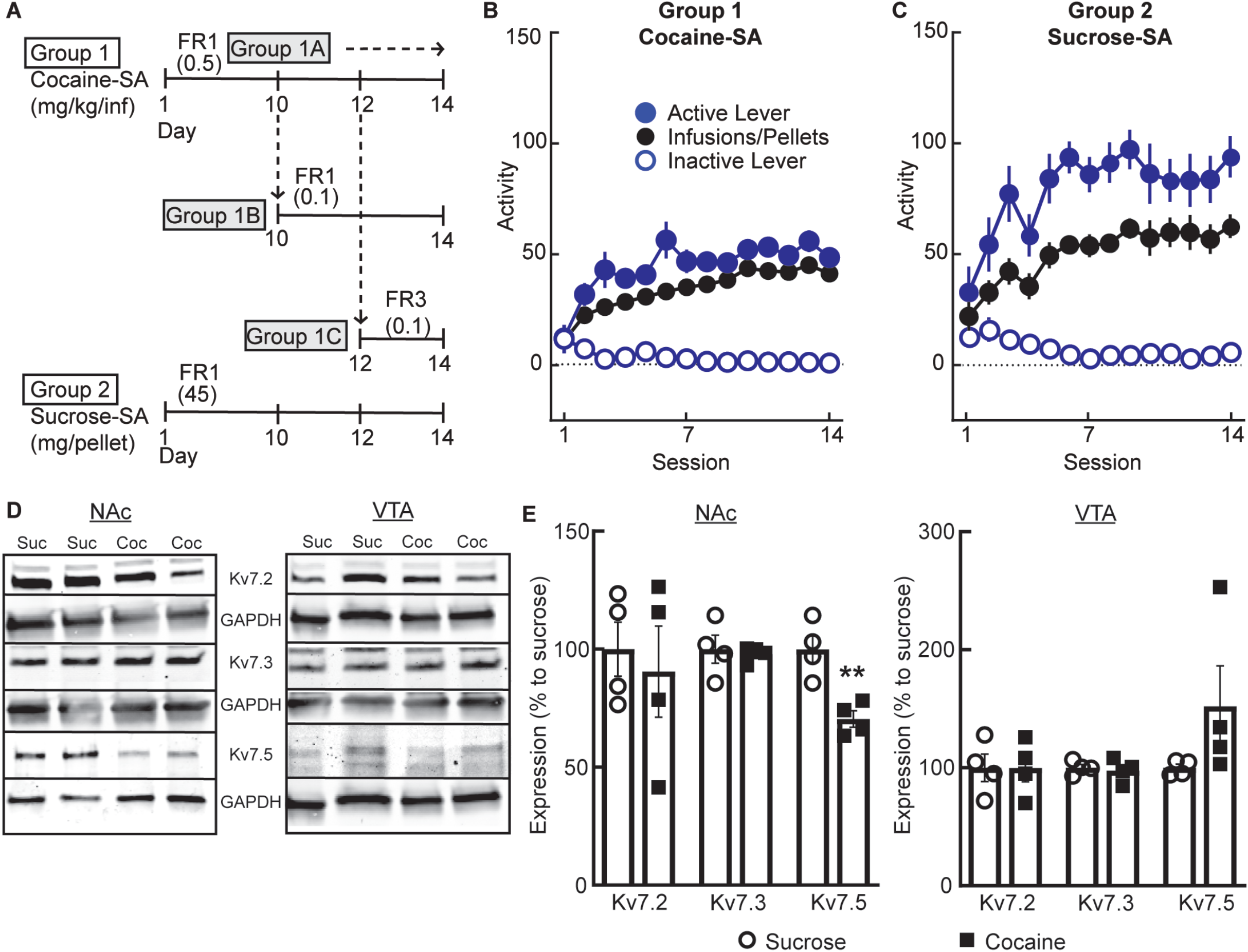
Training for self-administration of cocaine or sucrose and expression of the K_v_7 channel prior to retigabine testing. **A** Cohort design for rats learning to self-administer cocaine (cocaine-SA, Group-1; 0.5 mg/kg unit dose) or sucrose-SA (Group-2; 45 mg pellet). After d10-12 daily sessions, cocaine-SA rats remained in the same paradigm (Group-1A, 0.5 mg/kg/infusion; FR1) or transitioned to a different dose (Group-1B, 0.1 mg/kg/infusion; FR1) or dose and reinforcement schedule (Group-1C, 0.1 mg/kg/infusion; FR3). **B**,**C** Summary of behavioral responding when rats were presented with levers that resulted in the activation of a light and tone cue paired with **(B)** a cocaine infusion or **(C)** a sucrose pellet. **D**,**E** Western blots of NAc and VTA punch lysates showing K_v_7 subunit expression (7.2, 7.3, 7.5) normalized to GAPDH and expressed as a percentage of sucrose controls. **E** In the NAc, Kv7.5 expression decreased, whereas in the VTA, K_v_7 subunit expression did not differ between treatment groups. **p=0.0052 compared to sucrose-SA using an unpaired t-test

**Fig. 3.**
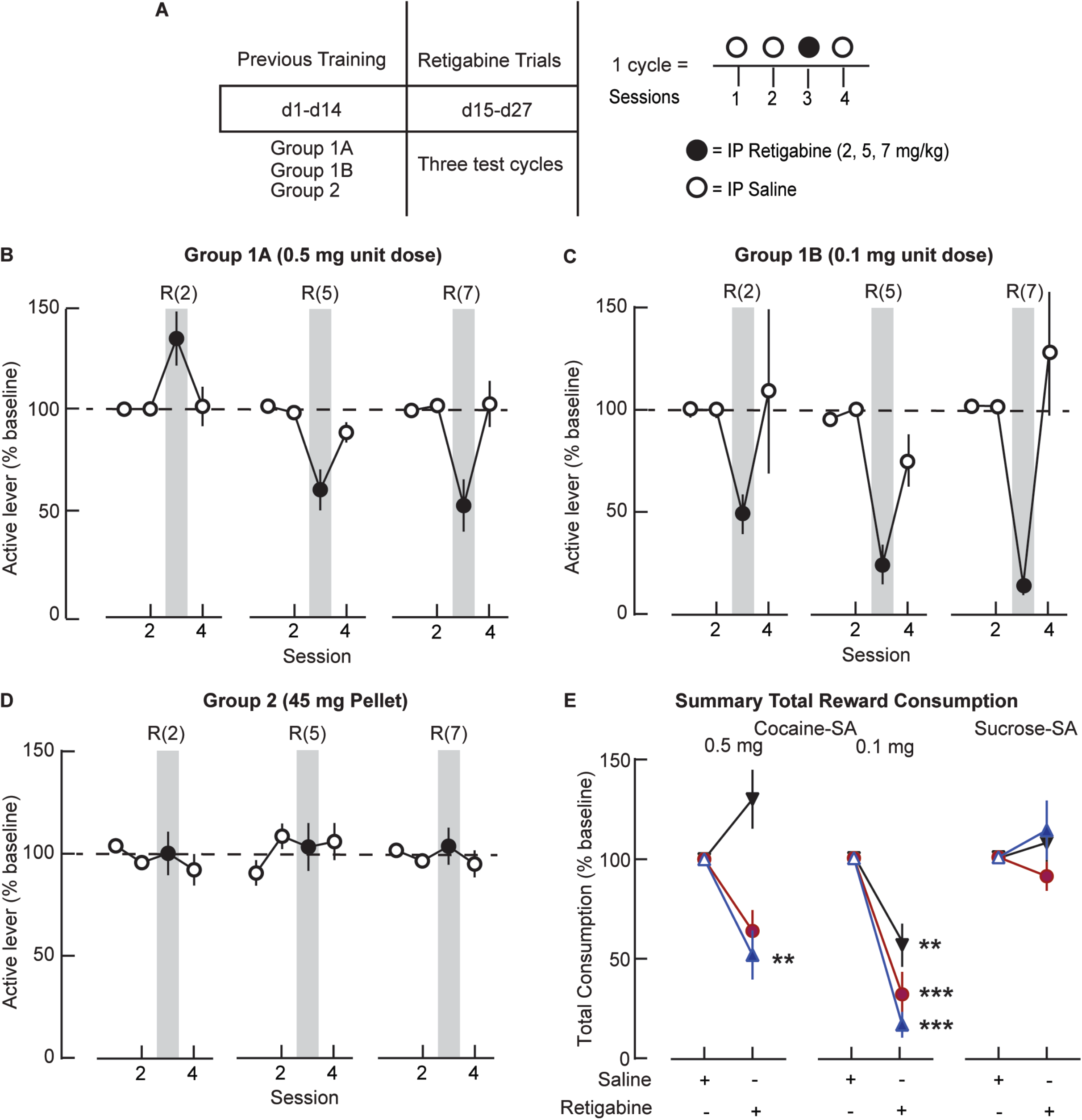
Retigabine reduces cocaine-but not sucrose-SA. **A** Schematic depicting retigabine testing on a FR1. Each group underwent 3 cycles (4 sessions per cycle) of testing with pretreatments with saline (1 ml/kg; Sessions-1,2,4) or increasing doses of retigabine (2, 5, 7 mg/kg, ip; Session-3) in a repeated-measure design administered 15 min before SA. **B**,**C**,**D** Summary data showing active lever responding in consecutive sessions (expressed as a percent of baseline) during a FR1 reinforcement schedule for cocaine at unit doses of **(B)** 0.5 mg **(C)** 0.1 mg or **(D)** sucrose (45 mg). Retigabine altered the active-lever responding for **(B**,**C)** cocaine at both unit doses, but not for **(D)** sucrose. *Gray bars* indicate retigabine pretreatment (R) at doses denoted by the numbers in parentheses (2, 5, 7 mg/kg, ip). **(E**) Summary of the total reward consumed after pretreatment with saline or different doses of retigabine (▾2 mg/kg, 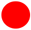 5 mg/kg, or 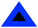 7 mg/kg). Significance compared to saline at the cocaine unit doses of 0.5 mg (7 mg/kg, **p=0.0133) and 0.1 mg (2 mg/kg, **p=0.0027; 5 and 7 mg/kg, ****p<0.0001) using a Šídák’s multiple comparison test

In addition to the 12-session FR1 training (above), animals in Group-1C (**Fig. 2A**) underwent two additional sessions of FR3 (each session 2h at 0.1 mg/kg/infusion), before transferring to cocaine-SA on a progressive-ratio (PR) schedule to test with retigabine. On the PR schedule (**Fig. 4A**), 3 groups of rats self-administered cocaine at one of 3 different unit doses (0.06, 0.1, or 0.25 mg/kg/infusion) with three days of FR3 between PR sessions (method adapted from ^21^. The number of active-lever responses required for successive reinforcers increased according to the published formula ^22^. The PR sessions ended after reaching a breakpoint or after 3 hours had elapsed, whichever occurred first.

**Fig. 4.**
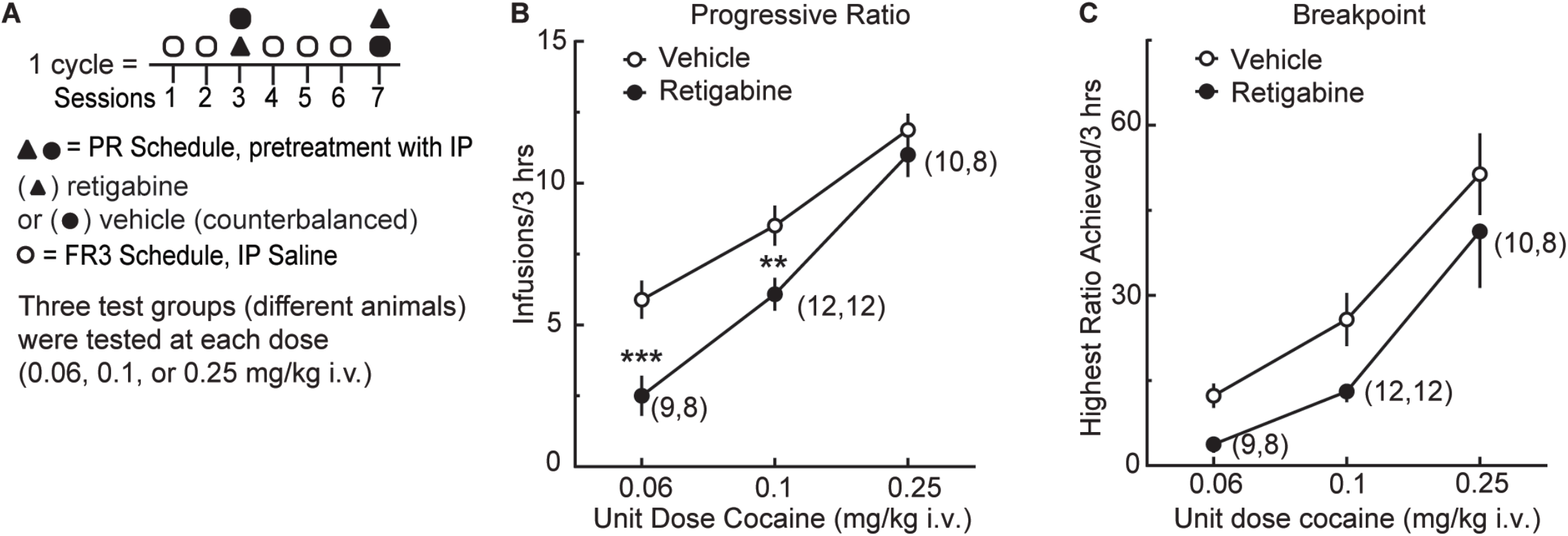
Retigabine reduces the motivation for cocaine in the progressive-ratio task. **A** Schematic depicting retigabine testing in a PR schedule of reinforcement. Group-1C rats received pretreatments with saline (1 ml/kg; Sessions-1,2,4-6) 15 min prior to session of cocaine-SA (0.1 mg unit dose) on the FR3 reinforcement schedule. In Sessions-3,-7, rats received vehicle (1ml/kg) or retigabine (5 mg/kg, IP) pretreatments 15 min before cocaine (0.06, 0.1, or 0.25 mg/kg) in a PR reinforcement schedule. Separate groups of rats were used for the different doses of cocaine. **B**,**C**, Effects of retigabine on (**B**) infusions and (**C**) breakpoints under a PR schedule of cocaine reinforcement at various doses. Numbers in paratheses denote the number of subjects. ***p=0.0002 (0.06 mg unit dose, infusions), **p=0.0021 (0.1 mg unit dose, infusions) compared to vehicle using a Šídák’s multiple comparison test; In (**B**), main effect, F_2,29_=12.71, p<0.0001; retigabine F_1,24_=10.63, p<0.0033; dose X retigabine F_2,24_=0.7867, p=0.4667

### Sucrose-SA

Naїve, ad libitum fed rats (n=12) were trained on a FR1 sucrose-SA paradigm (20s timeout post infusion) for 3 hours per session (**Fig. 2A**), 1 session per day, 6 days per week as previously described ^9^. The operant chambers and procedures were identical to the cocaine-SA described above except that training sessions paired the discrete cues with a sucrose pellet (45 mg, Bio-Serv). After acquisition of sucrose-SA, prior to retigabine, animals were habituated to (IP) pretreatment injections with a minimum of two days of saline treatments (1 ml/kg, IP) before retigabine and every day thereafter if not receiving retigabine (**Fig. 3A**).

### Drugs

Retigabine (Axon Medchem) was dissolved in saline with 10% Tween-80 (Sigma-Aldrich) at concentrations of 2, 5, or 7 mg/ml. Retigabine or vehicle pretreatments were administered 15 min before CPP or SA. Initial doses of retigabine were based on published work ^23^. Cocaine hydrochloride (Drug Supply Program of the National Institute on Drug Abuse) was prepared in sterile saline.

### Western blot

Using our published procedures ^9^, we prepared blots from a subset of Group-1A sucrose-SA (n=4) and cocaine-SA (n=4) rats. Approximately 10 ug of total proteins were loaded onto a SDS-PAGE gel and then transferred to nitrocellulose membranes and blocked at room temperature (1h). Primary antibodies for probing KCNQ2 (catalog #APC-050, Alomone Labs), KCNQ3 (catalog #APC-051, Alomone Labs), KCNQ5 (catalog #APC155, Alomone Labs), and GAPDH (catalog #MA1-16757, Invitrogen) were diluted in TBST with 5% BSA. The secondaries used were LiCor Goat anti-Rabbit IRDye800 (1:5000) and LiCor Goat anti-Mouse IRDye680 (1:5000). Images were captured using an Azure Sapphire imager, and blot densities analyzed using Azure Spot analysis software.

### Statistical analysis

All values are expressed as means ±SEM. Differences between groups were assessed using *t*-tests, one-way, or two-way repeated measures mixed-effects model. When significant main effects were obtained, appropriate post-hoc comparisons between groups were made using a Šídák’s multiple comparisons test. Significance was set at α = 0.05. All analyses were performed on Prism version 9.4.0 (GraphPad, La Jolla, CA).

## RESULTS

### Retigabine decreases cocaine CPP

Retigabine reduces dopamine neuron firing associated with psychostimulants ^18^, but its functional impact on cocaine-SA remains unknown. To evaluate this, we used a cocaine CPP (**Fig. 1A**) assay to first confirm a retigabine dose that effectively reduced the acquisition of a noncontingent reward behavior. For 5 days, we pretreated rats with retigabine (5 mg/kg, IP) or vehicle 15 min before daily counterbalanced treatments of cocaine or saline. On the post-test day, we tested the rats for chamber preference and found that in rats with a history of retigabine pretreatment, the time spent in the cocaine-paired chamber decreased relative to pretreatment with vehicle (t-test, t_20_=3.292; p=0.0036; **Fig. 1B**). These behavioral results identified a dose of retigabine sufficient to reduce noncontingent reward behavior.

### Training to self-administer cocaine or sucrose

To determine whether retigabine would also reduce the operant responding for a drug relative to a natural reward, we trained separate groups of male rats (**Fig. 2A)** to respond for cocaine (0.5 mg/kg/infusion; **Fig. 2B**) or sucrose pellets (**Fig. 2C**) on a FR1 schedule of reinforcement for at least 10 consecutive sessions. The rats readily learned to distinguish between active and inactive cocaine levers (lever: F_1,136_=243.0, p<0.0001; time: F_5.424,515.3_=2.078, p=0.0611; interaction F_13,1235_=5.965, p<0.0001) or sucrose (Lever: F_1,22_=86.72, p<0.0001; time: F_3.095,61.90_=2.674, p=0.0533; interaction F_13,260_=6.544, p<0.0001), indicating that the processing of reward reinforcement was intact. Because the effectiveness of retigabine may depend on the dose of cocaine or reinforcement schedule, we subdivided rats undergoing cocaine-SA (**Fig. 2A**) into three treatment groups to acclimate the animals to the different dose (0.5 mg/kg/infusion, FR1: Group-1A, n=29; 0.1 mg/kg/infusion, FR1: Groups-1B n=14) and/or reinforcement schedule (0.1 mg/kg/infusion, FR3: Group-1C, n=32). Comparing across cocaine and sucrose groups, we found no differences in the percentage of animals that reached the final criterion (Chi-square χ^2^_(3)_=1.584 [n=98], p=0.6630), indicating that the animals were ready for retigabine testing.

To determine whether the cocaine experiences altered the protein expression of K_v_7 channel subunits relative to sucrose, we sacrificed a subset of Group-1A sucrose-SA and cocaine-SA rats at 24hr after the final SA session for a Western blot analysis (**Fig. 2D**). We evaluated the main subunits of the neuronal K_v_7 family in the NAc and VTA tissue homogenates. In the NAc of cocaine-SA rats, the expression levels of K_v_7.5 were reduced (**Fig. 2E**; t_6_=4.277, p=0.0052), whereas the K_v_7.2 (t_6_=0.4209, p=0.6885) and K_v_7.3 (t_6_=0.3348, p=0.7491) were not. In contrast, in the VTA region, the K_v_7.2 (t_6_=0.0074, p=0.9943), the K_v_7.3, t_6_=0.4033, p=0.7007) and K_v_7.5 (t_6_=0.7127, p=0.5028) were similar between the drug and nondrug reward groups (**Fig. 2E**). Based on these results, we next determined whether the retigabine activation of inhibitory K_v_7 channels would reduce the operant responding for cocaine or sucrose.

### Retigabine testing with cocaine- or sucrose-SA

Using a 3-cycle within subject design (with 1 dose of retigabine per cycle; **Fig. 3A**), we compared the effects of saline with 3 doses of retigabine on the maintenance of reward SA behavior. In sessions 1 and 2, we injected rats with saline 15 min prior to SA sessions to habituate them to IP pretreatments. In session 3, we pretreated rats with retigabine (2, 5, or 7 mg/kg, IP) to test for changes in SA behavior. In cocaine-SA groups (**Fig. 3B**,**C)**, the substitution of retigabine for saline reduced active lever-pressing **(Fig. 3B** 0.5 mg/kg/infusion: treatment F_1,18_=8.218, *p*=0.0103; dose F_1.637,29.47_=7.579, *p*=0.0037; interaction F_1.921,17.29_=16.06, *p*=0.0001; **Fig. 3C** 0.1 mg/kg/infusion: treatment F_1,10_=55.79, *p*=0.0001; dose F_1.756,17.56_=1.190, *p*=0.3218; interaction F_1.917,9.585_=2.299, *p*=0.1539). Retigabine at 7 mg/kg, ip decreased cocaine responses by 2-fold at 0.5 mg/kg/infusion (**Fig. 3B)** and 6-fold at 0.1 mg/kg/ infusion (**Fig. 3C)**. In contrast, in sucrose rats (**Fig. 3D)** active lever-pressing was nearly unchanged in response to the same doses of retigabine (treatment F_1,11_=0.2843, *p*=0.6045; dose F_1.795,19.75_=1.560, *p*=0.2350; interaction F_1.771,19.48_=3.045, *p*=0.0759). We did not observe differences in IL responding for any of the groups. These results indicated that activation of K_v_7 channels could decrease the ability to maintain a SA pattern for cocaine, without altering similar responses for a natural reward.

To understand if the amount of reward consumed also decreased, we adjusted the change in the active lever responses for the total amount of reinforcer consumed in mg/kg and then expressed the data as a percentage of the baseline response to the preceding saline pretreatment (**Fig. 3E**). A two-way RM model with factors of pretreatment (saline or retigabine) and dose of retigabine revealed for the highest dose of cocaine (**Fig. 3E *left*)** a significant main effect of retigabine treatment (F_1,90_=5.795, *p*=0.0181) and dose (F_2,90_ = 11.40, *p*<0.0001), with an interaction between treatment and dose (F_2,90_=11.40, *p*<0.0001). The lower dose of cocaine (**Fig. 3E, *middle***) showed a similarly significant effect of retigabine (F_1,10_=71.71, *p*<0.0001), and dose (F_2,20_ = 6.295, *p*=0.0076), with an interaction between treatment and dose (F_2,10_=6.295, *p*=0.0170). However, sucrose animals (**Fig. 3E, *right***) showed no such response to retigabine (treatment: F_1,11_=0.2371, *p*=0.6359; dose: F_2,22_=1.287, *p*=0.2962; interaction: F_2,22_=1.287, *p*=0.2962). These results indicated that the retigabine responses differed between the cocaine- and sucrose-SA groups.

With the higher dose of cocaine (**Fig. 3E, *left*;** Group-1A: 0.5 mg/kg/infusion), responding on the active lever increased after 2 mg/kg retigabine and decreased after retigabine at 5 mg/kg and 7 mg/kg. Compared to saline pretreatment, only the reduction in cocaine consumption with the 7 mg/kg dose of retigabine differed significantly (Šídák’s multiple comparison test, t_180_=3.379, p=0.0133). On the contrary, with the lower dose of cocaine (**Fig. 3E, *middle***; Group-1B: 0.1 mg/kg/infusion), cocaine consumption decreased significantly relative to the saline baseline in all rats (2 mg/kg, t_20_=4.589, p=0.0027; 5 mg/kg, t_20_=6.541, p<0.0001; 7 mg/kg, t_20_=7.789, p<0.0001). The retigabine reduction was ∼3.5-fold greater with 7 mg/kg than 2 mg/kg retigabine (t_30_=4.938, p=0.004). Other changes with retigabine did not differ significantly from the saline baseline. These findings indicated dose-dependent actions of retigabine in the lower (Group-1B: 0.1 mg/kg/infusion) doses of cocaine.

Because the retigabine action did not generalize across reinforcers, we inspected the cumulative records of individual animals within the various reinforcer SA sessions (**Fig. 3B**,**C**,**D)** and made some general observations. First, we noted that the reductions related to retigabine in the distribution of the inter-infusion intervals for cocaine appeared dose-dependent. That is, in general for the cocaine sessions, the total number of infusions decreased as the dose of retigabine increased. However, retigabine failed to affect the distribution of inter-infusion intervals at either concentration of cocaine. Second, we observed that these changes were not evident in sucrose-SA rats. As an additional step, we evaluated responding on the inactive lever during pretreatment with saline for the three increasing doses of retigabine. A 2W ANOVA did not indicate significant differences related to retigabine in inactive lever responses within cocaine (Group-1A,B) or sucrose (Group-2) rats. Taken together, these results indicate that the pharmacological enhancement of K_v_7 channels with retigabine reduced cocaine administration, but the extent to which this occurred depended on both the dose of cocaine and retigabine. Lastly the effects of retigabine did not generalize to sucrose reward.

### Retigabine testing in a progressive-ratio task

The progressive-ratio (PR) schedule provides an index of reinforcement efficacy ^24^. As a final test, we used this approach to determine whether enhancing the activity of K_v_7 channels altered the motivation to earn a cocaine reward. Using Group-1C rats, we compared PR responses (**Fig. 4A**) after a pretreatment with saline (1 ml/kg, IP) versus retigabine (5 mg/kg, IP) in a repeated measure design. We used separate groups of animals to test the different doses of cocaine (0.06, 0.1, or 0.25 mg/kg/infusion). During PR tests, we found that responding for cocaine was dose-dependent, and that rats responded less for cocaine at lower doses (**Fig. 4B;** 2W RM mixed-effects model: F_2,29_=35.79, p<0.0001). There was also a main effect of treatment (**Fig. 4A**), indicating that when averaged between cocaine doses, rats exhibited lower infusion rates when treated with retigabine than when treated with vehicle (F_1,25_=29.01, p<0.0001). The difference between retigabine and vehicle was robust at a low dose of cocaine, but only modest at the highest dose of cocaine tested. There was a significant interaction (**Fig 4B)** of the dose of cocaine with retigabine (F_2,25_=3.627, p=0.0414). A post-hoc Šídák’s multiple comparison tests indicated that retigabine reduced infusions at the two lowest unit doses of cocaine (**Fig. 4B;** 0.06 mg unit dose: t_25_=4.696, p=0.0002; 0.1 mg unit dose: t_25_=3.857, p=0.0021; 0.25 mg unit dose: t_25_=0.9256, p=0.7421). Across the groups, the cocaine breakpoints increased as a function of the cocaine dose (main effect, F_2,29_=12.71, p<0.0001; **Fig. 4C**). However, the dose-response curve for cocaine was significantly decreased in response to retigabine (retigabine F_1,24_=10.63, p<0.0033; dose X retigabine F_2,24_=0.7867, p=0.4667; **Fig. 4C**). There were no main effects for the responses of the inactive lever (data not shown). In summary, retigabine significantly reduced cocaine intake over time and reduced incentive motivation for the drug. In contrast, at the doses tested, retigabine produced little measurable change in sucrose-SA.

## DISCUSSION

We identified K_v_7 channels as a regulator for the targeted control of chronic cocaine-SA. On a fixed-ratio reinforcement schedule, systemic treatments of retigabine dose-dependently reduced lever-pressing for cocaine but not the ‘natural’ reward, sucrose. These observations led us to pharmacologically evaluate K_v_7 channel activation during a progressive-ratio schedule of reinforcement, determining how retigabine shaped motivational responses at different doses of cocaine. Indeed, pretreatments with retigabine obstructed the motivational effects of cocaine at low to moderate doses. Therefore, we posit that, in combination with counseling and behavioral therapies, targeting K_v_7 channels may be a useful strategy for reducing the early stages of drug use and complement other treatments aimed at recovery during prolonged withdrawal ^25^. The present findings extend our previous research demonstrating that direct activation of cortical K_v_7 channels with retigabine reduced relapse-like behaviors ^9^ and align with recent accumbal work examining experimenter-delivered drug behaviors ^11^.

### Retigabine reduces cocaine CPP

To confirm an *in vivo* dose that would be effective in operant experiments, our research demonstrated a systemic dose of retigabine that blocked the acquisition of cocaine CPP. The reduction was greater than previously reported with flupirtine ^26^, a less selective structural precursor of retigabine that activates K_v_7.2/7.3 and other channels ^27^. The behavioral reduction with retigabine may reflect decreased dopamine transmission, as similar doses of retigabine reportedly decreased the enhancement of striatal and cortical dopamine levels associated with noncontingent administration of cocaine ^16^, which appear necessary for the acquisition of CPP ^28^.

### Retigabine actions in cocaine- or sucrose-SA behavior

We found that at lower doses of cocaine, K_v_7 channel activation reduced operant responses that had been learned and repeatedly reinforced in an FR paradigm. Although previous studies have not investigated this using operant behavior, retigabine reduced ethanol consumption in a two-bottle choice study ^29^ and alcohol phenotypes associated with K_v_7 channels ^30^. Although the underlying mechanisms of these and our studies remain to be determined, the critical role of phasic mesocortical dopamine signaling in reward-conditioning, learning, and SA behavior has been intensively investigated for many years ^31–34^. Dopamine depletion or receptor antagonism consistently impairs performance in instrumental tasks ^35–37^.

In addition to the examination of fixed-ratio responses, we confirmed the action of retigabine by measuring the motivational changes to obtain cocaine in a progressive-ratio reinforcement schedule ^38^. After confirming a previous progressive-ratio study showing that 0.06 to 0.25 unit doses supported cocaine-SA ^21^, we showed that retigabine was more effective at lowering PR responses in lower doses of cocaine. Earlier and more recent perspectives on progressive-ratio encompassing the perception of opportunity costs and decision making ^36,39^ agree with earlier work showing that the breakpoint is elevated when dopamine transmission is enhanced and attenuated when dopamine transmission is inhibited or depleted ^35,37,40–42^. Progressive-ratio responses for cocaine follow the same pattern ^24,43^.

Unlike cocaine, retigabine did not reduce sucrose-SA on a fixed-ratio schedule of reinforcement, which was relevant given that both rewards are associated with cue-induced seeking ^44^ and dopamine neuron activation ^45^. However, although dopamine release is time-locked to approach behavior for both cocaine and sucrose, dopamine levels quickly normalize during self-administration of sucrose, but not cocaine ^46,47^. Furthermore, unlike sucrose-SA, rats with a history of chronic cocaine-SA do not show the normal decrease in the firing of VTA dopamine neurons that occurs with learning ^48^. Furthermore, while cocaine-SA produces persistent LTP in VTA dopamine neurons, sucrose-SA does not ^49^. Although retigabine-LTP studies have not been reported in VTA dopamine neurons, in vivo administration of retigabine or flupirtine prevents hippocampal LTP and this effect was blocked by the selective K_v_7 channel antagonist XE991 ^50^. Furthermore, our previous work and others showed that direct cortical application of retigabine blocked the seeking for cocaine but not sucrose, the former of which requires VTA dopamine signaling in the prelimbic cortex ^51,52^.

### Expression of K_v_7 channel subtype after rewards

Our results confirmed earlier studies showing the expression of K_v_7 channel subunits in NAc and VTA ^53–55^. Unlike other reports linking decreases in brain K_v_7.2/ K_v_7.3 protein levels with alcohol withdrawal ^56^ or a depressive phenotype ^13,54,57^, we found no measurable differences between the drug and non-drug reward groups in the expression of K_v_7.2/K_v_7.3. Our observed reduction in NAc K_v_7.5 expression was interesting given that chronic SA of psychostimulants (including cocaine) and opiates upregulate cAMP-PKA signaling in NAc ^58,59^, and in culture systems, PKA phosphorylation robustly enhances K_v_7.5 containing channels ^60–62^. Chronic upregulation of cAMP-PKA signaling is associated with reward tolerance and dependence that drive SA ^59^. However, over time, chronic overactivation of cAMP-PKA signaling could produce homeostatic decreases in K_v_7.5 protein expression. How this would impact overall mesolimbic signaling is unclear, but in the hippocampus, reduced expression of inhibitory channels containing K_v_7.5 is associated with increased excitability ^63–65^.

## CONCLUSIONS

Our behavioral data align with published behavioral and electrophysiology work causally linking K_v_7 channel-mediated inhibition to reductions in dopamine neuron activity ^10^. Doses of retigabine similar to those used in our study reduced the frequency of VTA action potential firing and reduced psychostimulant-induced increases in dopamine release from the terminals ^15,18^. In hippocampal and VTA dopamine cells, retigabine more potently suppressed burst firing activity than basal neuron firing ^18,66^. Therefore, a reasonable scenario is that retigabine activation of K_v_7 channels attenuated excessive dopaminergic neurotransmission in the mesolimbic system, to reduce cocaine-SA.

Although our data provide a more inclusive evaluation of retigabine and SA is a well validated model with predictive value for medications that effectively treat cocaine dependence, the utility of retigabine for human substance abuse disorder is unclear. Whereas preclinical studies often use acute medication pretreatments to evaluate SA, most human studies instead assess cocaine craving or “high ^67^.” In controlled clinical trials, cocaine-SA appears difficult to diminish and may differ at different stages of the addiction cycle. Encouraging examples which decrease cocaine use include modafinil, baclofen and buprenorphine, although the latter two are limited by the issues of dose or enhancement of cocaine intoxication ^67^. Retigabine has an acceptable safety profile ^68^, and may prove useful as an adjunct therapy for psychosocial treatments to reduce maladaptive behavior. Although our results with retigabine in the rat sucrose-SA model were encouraging, it is unclear whether retigabine has unexpected adverse side effects in patients using drugs which could limit compliance. Taken together, our work emphasizes the potential value of future clinical trials with retigabine and further identification of its mechanisms relevant to drug self-administration.

## Contributions

Esteban S. Urena: Conceptualization, Methodology, Data Collection, Writing – original draft, Writing – Review & Editing. Cody C. Diezel: Conceptualization, Methodology, Data Collection. Mauricio Serna: Data Curation, Data Collection, Writing – original draft, Writing – Review & Editing. Grace Hala ’ufla: Data Collection. Kara R. Barber: Data collection, Data curation, Conceptualization, Methodology, Writing – original draft. Lisa Majuta: Data Collection, Methodology, Supervision, Writing – Review & Editing. Todd W. Vanderah: Writing – reviewing & editing. Arthur C. Riegel: Conceptualization, Writing – review & editing, Supervision

## Acknowledgments

The authors thank Dr. John M. Streicher for Western blot reagents, the Comprehensive Pain and Addiction-Center (CPA-C), and the Center of Excellence in Addiction Studies (CEAS) NIH/NIDA DA051255 for their support. The authors also thank the Vanderah lab for use of their CPP chambers.

## References

1. Juarez B, Han MH. Diversity of Dopaminergic Neural Circuits in Response to Drug Exposure. 2016;41(10):2424–2446. doi:10.1038/npp.2016.32

2. Kuhn BN, Kalivas PW, Bobadilla AC. Understanding Addiction Using Animal Models. Front Behav Neurosci. 2019;13:262. doi:10.3389/fnbeh.2019.00262

3. Shapiro MS, Roche JP, Kaftan EJ, Cruzblanca H, Mackie K, Hille B. Reconstitution of Muscarinic Modulation of the KCNQ2/KCNQ3 K+ Channels That Underlie the Neuronal M Current. J Neurosci. 2000;20(5):1710–1721. doi:10.1523/jneurosci.20-05-01710.2000

4. Wang HS, Pan Z, Shi W, et al. KCNQ2 and KCNQ3 Potassium Channel Subunits: Molecular Correlates of the M-Channel. Science. 1998;282(5395):1890–1893. doi:10.1126/science.282.5395.1890

5. Wickenden AD, Zou A, Wagoner PK, Jegla T. Characterization of KCNQ5/Q3 potassium channels expressed in mammalian cells. Brit J Pharmacol. 2001;132(2):381–384. doi:10.1038/sj.bjp.0703861

6. Soh H, Springer K, Doci K, Balsbaugh JL, Tzingounis AV. KCNQ2 and KCNQ5 form heteromeric channels independent of KCNQ3. Proc National Acad Sci. 2022;119(13):e2117640119. doi:10.1073/pnas.2117640119

7. Zhang J, Carver CM, Choveau FS, Shapiro MS. Clustering and Functional Coupling of Diverse Ion Channels and Signaling Proteins Revealed by Super-resolution STORM Microscopy in Neurons. Neuron. 2016;92(2):461–478. doi:10.1016/j.neuron.2016.09.014

8. M. S M, M. M, J. M S, A. B D, P. D. Molecular correlates of the M-current in cultured rat hippocampal neurons. J Physiol (Lond). 2002;544:29–37. doi:10.1113/jphysiol.2002.028571

9. J. PC, C. B W, P. G, et al. Restoration of Kv7 Channel-Mediated Inhibition Reduces Cued-Reinstatement of Cocaine Seeking. J Neurosci. 2018;38(17):4212–4229. doi:10.1523/jneurosci.2767-17.2018

10. Koyama S, Appel SB. Characterization of M-current in ventral tegmental area dopamine neurons. Journal of neurophysiology. 2006;96(2):535–543. doi:10.1152/jn.00574.2005

11. Tsuboi D, Otsuka T, Shimomura T, et al. Dopamine drives neuronal excitability via KCNQ channel phosphorylation for reward behavior. Cell Reports. 2022;40(10):111309. doi:10.1016/j.celrep.2022.111309

12. Vigil FA, Carver CM, Shapiro MS. Pharmacological Manipulation of K v 7 Channels as a New Therapeutic Tool for Multiple Brain Disorders. Front Physiol. 2020;11:688. doi:10.3389/fphys.2020.00688

13. Friedman AK, Juarez B, Ku SM, et al. KCNQ channel openers reverse depressive symptoms via an active resilience mechanism. Nat Commun. 2016;7(1):11671. doi:10.1038/ncomms11671

14. Jensen MM, Lange SC, Thomsen MS, Hansen HH, Mikkelsen JD. The Pharmacological Effect of Positive KCNQ (Kv7) Modulators on Dopamine Release from Striatal Slices. Basic Clin Pharmacol. 2011;109(5):339–342. doi:10.1111/j.1742-7843.2011.00730.x

15. Hansen HH, Ebbesen C, Mathiesen C, et al. The KCNQ Channel Opener Retigabine Inhibits the Activity of Mesencephalic Dopaminergic Systems of the Rat. J Pharmacol Exp Ther. 2006;318(3):1006–1019. doi:10.1124/jpet.106.106757

16. Hansen HH, Andreasen JT, Weikop P, Mirza N, Scheel-Krüger J, Mikkelsen JD. The neuronal KCNQ channel opener retigabine inhibits locomotor activity and reduces forebrain excitatory responses to the psychostimulants cocaine, methylphenidate and phencyclidine. European journal of pharmacology. 2007;570(1-3):77–88. doi:10.1016/j.ejphar.2007.05.029

17. Martire M, D‘Amico M, Panza E, et al. Involvement of KCNQ2 subunits in [3H]dopamine release triggered by depolarization and pre‐synaptic muscarinic receptor activation from rat striatal synaptosomes. J Neurochem. 2007;102(1):179–193. doi:10.1111/j.1471-4159.2007.04562.x

18. Sotty F, Damgaard T, Montezinho LP, et al. Antipsychotic-Like Effect of Retigabine [N-(2-Amino-4-(fluorobenzylamino)-phenyl)carbamic Acid Ester], a KCNQ Potassium Channel Opener, via Modulation of Mesolimbic Dopaminergic Neurotransmission. J Pharmacol Exp Ther. 2009;328(3):951–962. doi:10.1124/jpet.108.146944

19. Grenald SA, Largent-Milnes TM, Vanderah TW. Animal models for opioid addiction drug discovery. Expert Opin Drug Dis. 2014;9(11):1345–1354. doi:10.1517/17460441.2014.966076

20. Brown L, Gutherz S, Kulick C, Soper C, Kondratyev A, Forcelli PA. Profile of retigabine‐induced neuronal apoptosis in the developing rat brain. Epilepsia. 2016;57(4):660–670. doi:10.1111/epi.13335

21. Allain F, Bouayad-Gervais K, Samaha AN. High and escalating levels of cocaine intake are dissociable from subsequent incentive motivation for the drug in rats. Psychopharmacology. 2018;235(1):317–328. doi:10.1007/s00213-017-4773-8

22. McGregor A, Lacosta S, Roberts DCS. L-Tryptophan decreases the breaking point under a progressive ratio schedule of intravenous cocaine reinforcement in the rat. Pharmacol Biochem Be. 1993;44(3):651–655. doi:10.1016/0091-3057(93)90181-r

23. Blackburn‐Munro G, Dalby‐Brown W, Mirza NR, Mikkelsen JD, Blackburn‐Munro RE. Retigabine: Chemical Synthesis to Clinical Application. Cns Drug Rev. 2005;11(1):1–20. doi:10.1111/j.1527-3458.2005.tb00033.x

24. Stafford D, LeSage MG, Glowa JR. Progressive-ratio schedules of drug delivery in the analysis of drug self-administration: a review. Psychopharmacology. 1998;139(3):169–184. doi:10.1007/s002130050702

25. Chiu AS, Kang MC, Sanchez LLH, et al. Preclinical evidence to support repurposing everolimus for craving reduction during protracted drug withdrawal. Neuropsychopharmacol. 2021;46(12):2090–2100. doi:10.1038/s41386-021-01064-9

26. Mooney J, Rawls SM. KCNQ2 & sol;3 channel agonist flupirtine reduces cocaine place preference in rats. Behav Pharmacol. 2017;28(5):405–407. doi:10.1097/fbp.0000000000000287

27. Kornhuber J, Bleich S, Wiltfang J, Maler M, Parsons CG. Flupirtine shows functional NMDA receptor antagonism by enhancing Mg2+ block via activation of voltage independent potassium channels. J Neural Transmission. 1999;106(9-10):857–867. doi:10.1007/s007020050206

28. Kawahara Y, Ohnishi YN, Ohnishi YH, Kawahara H, Nishi A. Distinct Role of Dopamine in the PFC and NAc During Exposure to Cocaine-Associated Cues. Int J Neuropsychoph. 2021;24(12):988–1001. doi:10.1093/ijnp/pyab067

29. Knapp CM, O‘Malley M, Datta S, Ciraulo DA. The Kv7 potassium channel activator retigabine decreases alcohol consumption in rats. Am J Drug Alcohol Abus. 2014;40(3):244–250. doi:10.3109/00952990.2014.892951

30. McGuier NS, Rinker JA, Cannady R, et al. Identification and validation of midbrain Kcnq4 regulation of heavy alcohol consumption in rodents. Neuropharmacology. 2018;138:10–19. doi:10.1016/j.neuropharm.2018.05.020

31. Fischbach-Weiss S, Reese RM, Janak PH. Inhibiting Mesolimbic Dopamine Neurons Reduces the Initiation and Maintenance of Instrumental Responding. Neuroscience. 2018;372:306–315. doi:10.1016/j.neuroscience.2017.12.003

32. Aberman JE, Salamone JD. Nucleus accumbens dopamine depletions make rats more sensitive to high ratio requirements but do not impair primary food reinforcement. Neuroscience. 1999;92(2):545–552. doi:10.1016/s0306-4522(99)00004-4

33. Aberman JE, Ward SJ, Salamone JD. Effects of Dopamine Antagonists and Accumbens Dopamine Depletions on Time-Constrained Progressive-Ratio Performance. Pharmacol Biochem Be. 1998;61(4):341–348. doi:10.1016/s0091-3057(98)00112-9

34. Mingote S, Weber SM, Ishiwari K, Correa M, Salamone JD. Ratio and time requirements on operant schedules: effort‐related effects of nucleus accumbens dopamine depletions. Eur J Neurosci. 2005;21(6):1749–1757. doi:10.1111/j.1460-9568.2005.03972.x

35. Nicola SM. The Flexible Approach Hypothesis: Unification of Effort and Cue-Responding Hypotheses for the Role of Nucleus Accumbens Dopamine in the Activation of Reward-Seeking Behavior. J Neurosci. 2010;30(49):16585–16600. doi:10.1523/jneurosci.3958-10.2010

36. Salamone JD, Correa M. The mysterious motivational functions of mesolimbic dopamine. Neuron. 2012;76(3):470–485. doi:10.1016/j.neuron.2012.10.021

37. Salamone JD, Wisniecki A, Carlson BB, Correa M. Nucleus accumbens dopamine depletions make animals highly sensitive to high fixed ratio requirements but do not impair primary food reinforcement. Neuroscience. 2001;105(4):863–870. doi:10.1016/s0306-4522(01)00249-4

38. Richardson NR, Roberts DCS. Progressive ratio schedules in drug self-administration studies in rats: a method to evaluate reinforcing efficacy. J Neurosci Meth. 1996;66(1):1–11. doi:10.1016/0165-0270(95)00153-0

39. Salamone JD. Commentary: WILL THE LAST PERSON WHO USES THE TERM ‘REWARD ‘ PLEASE TURN OUT THE LIGHTS? COMMENTS ON PROCESSES RELATED TO REINFORCEMENT, LEARNING, MOTIVATION AND EFFORT. Addict Biol. 2006;11(1):43–44. doi:10.1111/j.1369-1600.2006.00011.x

40. Hamill S, Trevitt JT, Nowend KL, Carlson BB, Salamone JD. Nucleus Accumbens Dopamine Depletions and Time-Constrained Progressive Ratio Performance Effects of Different Ratio Requirements. Pharmacol Biochem Be. 1999;64(1):21–27. doi:10.1016/s0091-3057(99)00092-1

41. Zhang M, Balmadrid C, Kelley AE. Nucleus Accumbens Opioid, GABAergic, and Dopaminergic Modulation of Palatable Food Motivation: Contrasting Effects Revealed by a Progressive Ratio Study in the Rat. Behav Neurosci. 2003;117(2):202–211. doi:10.1037/0735-7044.117.2.202

42. Baldo BA, Jain K, Veraldi L, Koob GF, Markou A. A Dopamine D1 Agonist Elevates Self-Stimulation Thresholds Comparison to Other Dopamine-Selective Drugs. Pharmacol Biochem Be. 1999;62(4):659–672. doi:10.1016/s0091-3057(98)00206-8

43. Roberts DCS, Corcoran ME, Fibiger HC. On the role of ascending catecholaminergic systems in intravenous self-administration of cocaine. Pharmacol Biochem Be. 1977;6(6):615–620. doi:10.1016/0091-3057(77)90084-3

44. Bobadilla AC, Dereschewitz E, Vaccaro L, Heinsbroek JA, Scofield MD, Kalivas PW. Cocaine and sucrose rewards recruit different seeking ensembles in the nucleus accumbens core. Mol Psychiatr. Published online 2020:1-14. doi:10.1038/s41380-020-00888-z

45. Alsio J, Nordenankar K, Arvidsson E, et al. Enhanced Sucrose and Cocaine Self-Administration and Cue-Induced Drug Seeking after Loss of VGLUT2 in Midbrain Dopamine Neurons in Mice. J Neurosci. 2011;31(35):12593–12603. doi:10.1523/jneurosci.2397-11.2011

46. Roitman MF, Stuber GD, Phillips PEM, Wightman RM, Carelli RM. Dopamine operates as a subsecond modulator of food seeking. The Journal of neuroscience : the official journal of the Society for Neuroscience. 2004;24(6):1265–1271. doi:10.1523/jneurosci.3823-03.2004

47. Phillips PEM, Stuber GD, Heien MLAV, Wightman RM, Carelli RM. Subsecond dopamine release promotes cocaine seeking. NATURE-LONDON-. 2003;422(6932):614–618. doi:10.1038/nature01476

48. Takahashi YK, Stalnaker TA, Marrero-Garcia Y, Rada RM, Schoenbaum G. Expectancy-Related Changes in Dopaminergic Error Signals Are Impaired by Cocaine Self-Administration. Neuron. 2019;101(2):294–306.e3. doi:10.1016/j.neuron.2018.11.025

49. Chen BT, Bowers MS, Martin M, et al. Cocaine but not natural reward self-administration nor passive cocaine infusion produces persistent LTP in the VTA. Neuron. 2008;59(2):288–297. doi:10.1016/j.neuron.2008.05.024

50. Li C, Huang P, Lu Q, Zhou M, Guo L, Xu X. KCNQ/Kv7 channel activator flupirtine protects against acute stress-induced impairments of spatial memory retrieval and hippocampal LTP in rats. Neuroscience. 2014;280:19–30. doi:10.1016/j.neuroscience.2014.09.009

51. James MH, McGlinchey EM, Vattikonda A, Mahler SV, Aston-Jones G. Cued Reinstatement of Cocaine but Not Sucrose Seeking Is Dependent on Dopamine Signaling in Prelimbic Cortex and Is Associated with Recruitment of Prelimbic Neurons That Project to Contralateral Nucleus Accumbens Core. Int J Neuropsychoph. 2017;21(1):89–94. doi:10.1093/ijnp/pyx107

52. C. B W, V. M S, B. H, S. AJ G, C. R A. Dopamine terminals from the ventral tegmental area gate intrinsic inhibition in the prefrontal cortex. Physiological Reports. 2017;5(6):e13198. doi:10.14814/phy2.13198

53. Faruk MdO, Tsuboi D, Yamahashi Y, et al. Muscarinic signaling regulates voltage‐gated potassium channel KCNQ2 phosphorylation in the nucleus accumbens via protein kinase C for aversive learning. J Neurochem. Published online 2021. doi:10.1111/jnc.15555

54. Li L, Sun H, Ding J, et al. Selective targeting of M‐type potassium Kv7.4 channels demonstrates their key role in the regulation of dopaminergic neuronal excitability and depression‐like behaviour. Brit J Pharmacol. 2017;174(23):4277–4294. doi:10.1111/bph.14026

55. Su M, Li L, Wang J, et al. Kv7.4 Channel Contribute to Projection-Specific Auto-Inhibition of Dopamine Neurons in the Ventral Tegmental Area. Front Cell Neurosci. 2019;13:557. doi:10.3389/fncel.2019.00557

56. Kang S, Li J, Zuo W, et al. Ethanol Withdrawal Drives Anxiety-Related Behaviors by Reducing M-type Potassium Channel Activity in the Lateral Habenula. Neuropsychopharmacol. 2017;42(9):1813–1824. doi:10.1038/npp.2017.68

57. Krishnan V, Han MH, Graham DL, et al. Molecular adaptations underlying susceptibility and resistance to social defeat in brain reward regions. Cell. 2007;131(2):391–404. doi:10.1016/j.cell.2007.09.018

58. Lu L, Grimm JW, Shaham Y, Hope BT. Molecular neuroadaptations in the accumbens and ventral tegmental area during the first 90 days of forced abstinence from cocaine self‐administration in rats. J Neurochem. 2003;85(6):1604–1613. doi:10.1046/j.1471-4159.2003.01824.x

59. Nestler EJ. Reflections on: “A general role for adaptations in G-Proteins and the cyclic AMP system in mediating the chronic actions of morphine and cocaine on neuronal function.” Brain Res. 2016;1645:71–74. doi:10.1016/j.brainres.2015.12.039

60. Brueggemann LI, Cribbs LL, Schwartz J, Wang M, Kouta A, Byron KL. Mechanisms of PKA-Dependent Potentiation of Kv7.5 Channel Activity in Human Airway Smooth Muscle Cells. Int J Mol Sci. 2018;19(8):2223. doi:10.3390/ijms19082223

61. Mani BK, Robakowski C, Brueggemann LI, et al. Kv7.5 Potassium Channel Subunits Are the Primary Targets for PKA-Dependent Enhancement of Vascular Smooth Muscle Kv7 Currents. Mol Pharmacol. 2016;89(3):323–334. doi:10.1124/mol.115.101758

62. Horst J van der, Greenwood IA, Jepps TA. Cyclic AMP-Dependent Regulation of Kv7 Voltage-Gated Potassium Channels. Front Physiol. 2020;11:727. doi:10.3389/fphys.2020.00727

63. Tzingounis AV, Heidenreich M, Kharkovets T, et al. The KCNQ5 potassium channel mediates a component of the afterhyperpolarization current in mouse hippocampus. PNAS. 2010;107(22):10232–10237. doi:10.1073/pnas.1004644107

64. E. YN, A. M, N. S, et al. Localization of KCNQ5 in the normal and epileptic human temporal neocortex and hippocampal formation. Neuroscience. 2003;120:353–364. doi:10.1016/s0306-4522(03)00321-x

65. Lehman A, Thouta S, Mancini GMS, et al. Loss-of-Function and Gain-of-Function Mutations in KCNQ5 Cause Intellectual Disability or Epileptic Encephalopathy. Am J Hum Genetics. 2017;101(1):65–74. doi:10.1016/j.ajhg.2017.05.016

66. Cuiyong Y, Yoel Y. KCNQ/M Channels Control Spike Afterdepolarization and Burst Generation in Hippocampal Neurons. Journal of Neuroscience. 2004;24:4614–4624. doi:10.1523/jneurosci.0765-04.2004

67. Haney M, Spealman R. Controversies in translational research: drug self-administration. Psychopharmacology. 2008;199(3):403–419. doi:10.1007/s00213-008-1079-x

68. Faulkner MA, Burke RA. Safety profile of two novel antiepileptic agents approved for the treatment of refractory partial seizures: ezogabine (retigabine) and perampanel. Expert Opin Drug Saf. 2013;12(6):847–855. doi:10.1517/14740338.2013.823399

